# Layer-Weighted Source-Anchored Transfer Learning for Multi-Omics Survival Prediction and Risk Stratification in Younger Breast Cancer Patients

**DOI:** 10.64898/2026.11.01.763494

**Authors:** Wanru Guo, Binbin Wang, Curtis Tatsuoka

**Affiliations:** Department of Epidemiology and Public Health, Division of Biostatistics and Bioinformatics, University of Maryland–Baltimore, Baltimore, Maryland, USA; University of Maryland Greenebaum Comprehensive Cancer Center, Baltimore, Maryland, USA

## Abstract

**Background:** Younger breast cancer patients (<50 years) often present with more aggressive disease and distinct molecular characteristics, yet predictive models for this group are underpowered due to small sample sizes. Most existing models are built on older populations and may not translate directly, particularly across multiple omics data types. To address this, we developed *Layered Source-Anchored Transfer Elastic Net (LSAT-EN)*, which adaptively borrows from the older source cohort while allowing borrowing strength to vary across omics layers according to their similarity between source and target cohorts, while performing EN variable selection.

**Methods:** Patients aged ≥ 60, ≥55, and ≥50 years were used as the source to improve prediction in patients aged <50 years. LSAT-EN were evaluated in group-correlated simulations and in TCGA-BRCA integrating 6 omics layers. Binary survival at 2–4 years was assessed by AUC; censored survival by Harrell’s C-index and 2–5-year inverse probability of censoring weighted (IPCW) AUC.

**Results:** In simulation, LSAT-EN achieved the highest AUC under moderate source-target heterogeneity (*σ*=0.25; 0.747–0.802), outperforming GAS-EN (0.735–0.767), overall EN (0.683–0.710), individual EN (0.646–0.767), and glmtrans EN (0.640–0.733) across target training sample sizes (nT). Its advantage was greatest under higher source-target heterogeneity (*σ*=4), particularly with small nT, where LSAT-EN achieved AUCs of 0.689–0.789 compared with 0.590–0.771 across competing methods. In TCGA-BRCA data, LSAT-EN improved discrimination relative to overall and individual EN in all binary and Cox models across all time horizons and source age groups. LSAT-EN was robust in stratifying high and low risk groups across molecular and clinical subgroups, while LSATEN-derived risk scores captured breast cancer-relevant pathway in survival models across time horizons, along with identifying most important miRNA signals.

**Conclusion:** LSAT-EN is a powerful TL method for improving prediction and risk stratification for younger breast cancer patients; The framework may support prediction in other underrepresented populations with limited target-cohort data to advance cancer disparity ML research.

## 1 Introduction

Breast cancer is the most commonly diagnosed cancer and the second leading cause of cancer death among women in the United States. Women younger than 50 accounted for 16% of estimated new invasive cases in 2024, yet incidence rose faster in this group during 2012–2021 than among women aged 50 or older (1.4% versus 0.7% annually). American Cancer Society (2024). Evidence from women diagnosed *before 40* also suggests that age can matter after diagnosis: Franco et al. (2023) reported more basal-like and HER2-positive tumors and a 1.52-fold higher disease-specific death hazard relative to women aged 40–60, while a (Raj-Kumar et al., 2025) identified differences in proliferation, hormone-response, and immune pathways. These findings motivate survival prediction that accounts for both the smaller younger cohort and potential differences in tumor biology.

Existing approaches offer two imperfect choices. An overall model pools patients across ages, as in Wang et al. (2022) SEER-based survival nomogram; an independent model uses only patients in the group of interest. Kang et al. (2024) fitted separate models for younger and older patients, while Gong et al. (2018) developed nomograms specifically for women under 40. Pooling provides more training data but may obscure age-specific associations; independent fitting avoids that assumption but has fewer observations and outcomes to learn from. Multi-omics prognosis has also been studied: SALMON (Huang et al., 2019) combined mRNA, miRNA, copy-number and mutation burdens, and clinical variables, but did not transfer a model between age groups.

Transfer learning offers a way to use a larger source cohort while adapting predictions to a smaller target cohort (Pan & Yang, 2010; Gao & Cui, 2020). In breast cancer, Gu et al. (2023) applied it to risk prediction in underrepresented populations, Wagner et al. (2026) adapted an imaging model from detection to risk prediction, and Pilgram et al. (2026) fine-tuned a survival prognostic tool. Multi-omics transfer has been studied across racial groups for progression-free interval in the MOTLAB preprint (Baek et al., 2025) and across cancer types, including breast cancer, in a (Wen & Li, 2025). To our knowledge, these studies do not evaluate older-to-younger transfer for multi-omics breast cancer survival prediction. That is the setting addressed here.

To address the limited outcome data available for younger patients with breast cancer, we propose LSAT-EN (Layer-specific source-anchored transfer learning with elastic-net regularization). We first fit a model in older patients (≥60/55/50 years) to estimate source coefficients, then fit a model in younger patients (<50 years) using those coefficients as an anchor. Unlike conventional source-anchored transfer learning, LSAT-EN tunes the strength of borrowing separately for each omics layer, alongside the elastic-net penalty. This allows not only prediction, but also variable selection/biomarker discovery as well as coefficient interpretation at the same time. We apply this procedure to logistic models for fixed-horizon binary outcomes and Cox models for censored survival outcomes.

Across the evaluated binary and survival analyses, LSAT-EN improved prediction in the younger target cohort relative to pooled, target-only, and conventional transfer-learning comparators in the reported settings, was robust in risk stratification across molecular and clinical subgroups, while LSAT EN-derived risk scores captured breast cancer-relevant pathways and miRNA signals. These findings support layer-specific borrowing as a way to use information from older patients while allowing prognostic associations to differ in younger patients.

## 2 Methods

### 2.1 Layer-specific source-anchored transfer learning with elastic-net regularization (LSAT-EN)

LSAT-EN transfers prognostic information from a source cohort to a target cohort for fixed-horizon binary and censored time-to-event prediction (Fig. 1). In each repetition, target patients undergo a stratified 70% training/30% testing split. Within each training partition, modality-specific quality filtering is followed by median imputation and feature standardization. Layer-wise SIS ranks eligible features separately within each omics layer using source and target-training outcomes. Screening budgets are selected through inner five-fold cross-validation from prespecified candidate counts: 250, 500, or 1,000 for mRNA and gene-level methylation; 50, 100, or 200 for miRNA; 100, 200, or 400 for CNV; 50, 100, or all eligible features for protein; and 25, 50, or 100 for mutation. Candidate counts are capped at the number of eligible features available within each training fold. Preprocessing and screening are repeated within the inner folds used to select the screening budgets and tune *λ*, *α*, and the layer-specific *τ_g_*. Source-only elastic-net models provide endpoint-specific coefficient anchors, re-estimated for each selected panel and preprocessing scale and held fixed during the corresponding target-model fits. Following tuning, the pipeline is refitted using all target-training data and the corresponding source anchor, then evaluated on the untouched target-test patients. Feature panels may therefore vary across repetitions. (Fan and Lv, 2008)

**Figure 1.**
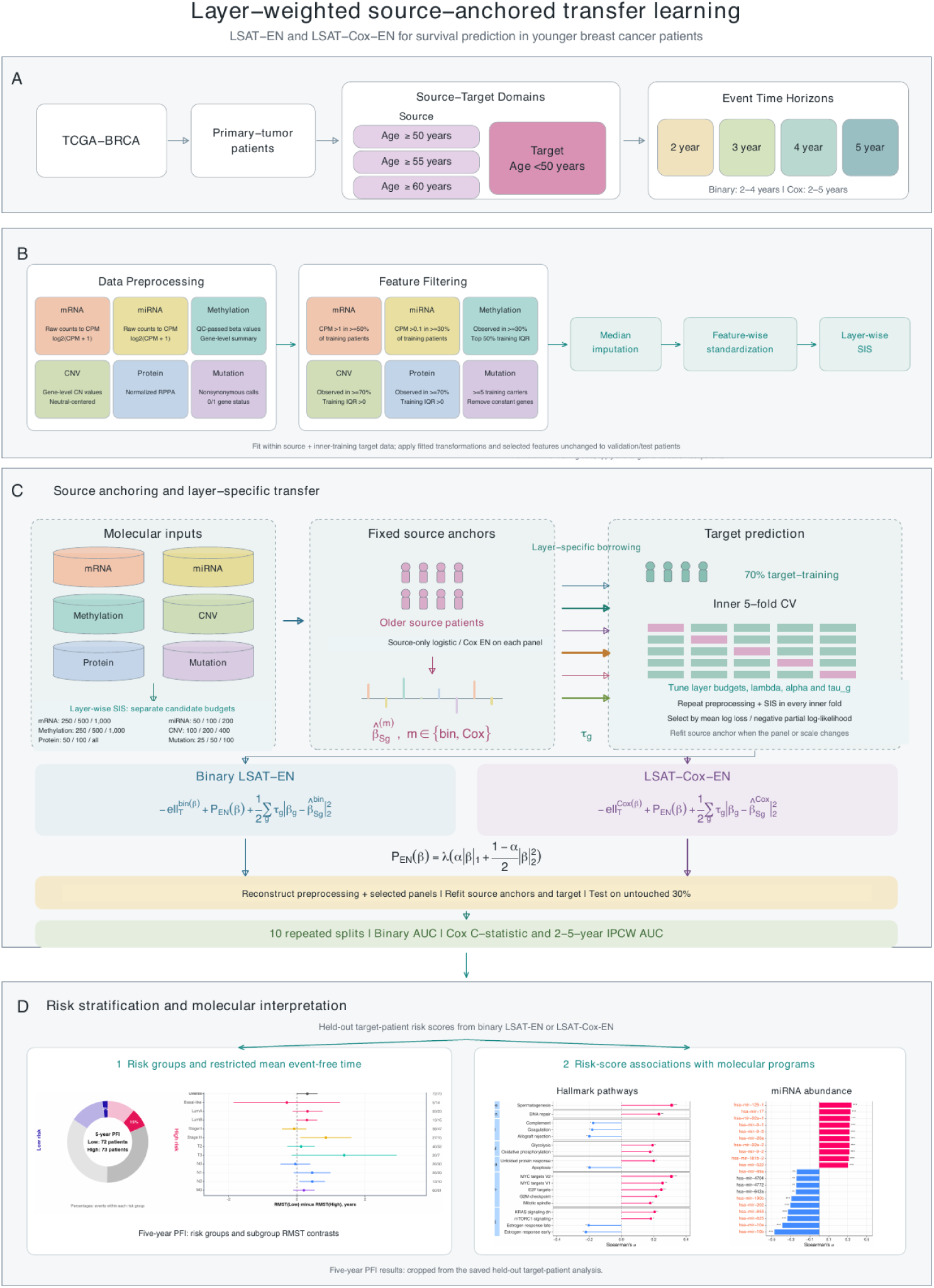
Workflow of LSAT-EN in prediction and risk stratification for younger BC patients. **A)** TCGA-BRCA primary-tumor patients define older source cohorts aged ≥60, ≥55, or ≥50 years and a younger target cohort aged <50 years, with binary survival prediction at 2–4 years and time-to-event evaluation at 2–5 years. **B)** Six molecular layers—mRNA, miRNA, DNA methylation, copy-number variation, protein abundance, and somatic mutations—are matched across patients and undergo modality-specific preprocessing and quality filtering, followed by training-fitted median imputation and feature-wise standardization. **C)** The workflow comprises five steps: 1) layer-wise SIS using source and target-training data to construct candidate panels with layer-specific screening budgets; 2) fitting source-only elastic-net models to obtain endpoint-specific coefficient anchors, held fixed during corresponding target-model fits; 3) selecting screening budgets and *λ*, *α*, and *τ* g through inner five-fold cross-validation within the 70% target-training subset, repeating preprocessing and screening within each fold and re-estimating anchors when the panel or scale changes; 4) reconstructing the selected pipeline using source and all target-training data, refitting source anchors and then target coefficients; and 5) evaluating the fitted pipeline on the untouched 30% target-test subset and reporting mean performance across 10 repeated splits. Binary evaluation uses AUC; time-to-event evaluation uses Harrell’s C-statistic and IPCW AUC. D) Held-out target-patient risk scores support risk grouping and Kaplan–Meier-based restricted mean event-free time comparisons, overall and within eligible clinical subgroups (left panel), alongside Spearman correlation analyses with Hallmark pathway activity and miRNA abundance (right panel).

#### 2.1.1 Fixed-horizon binary model

At horizon h, the binary response is 1 for a patient known to survive through h and 0 for death before h. Patients censored before h without a known outcome are excluded from that horizon’s binary analysis. With an unpenalized intercept, binary LSAT-EN minimizes the following target-training objective:

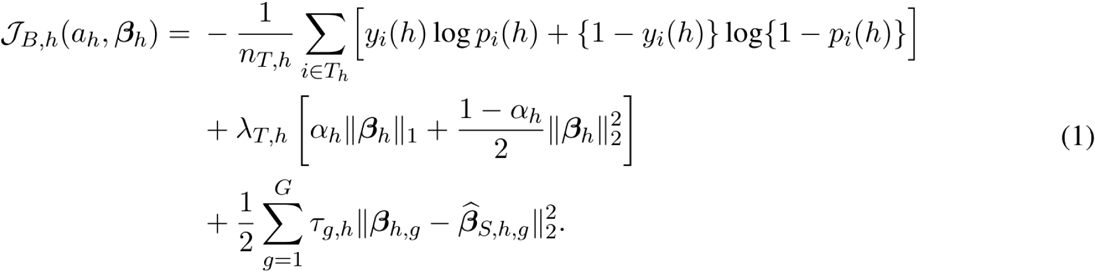

#### 2.1.2 Cox time-to-event model

The Cox analysis retains observed follow-up times and censoring indicators. Let the event indicator identify observed deaths and let the risk set contain patients still under observation at each event time. Cox LSAT-EN minimizes:

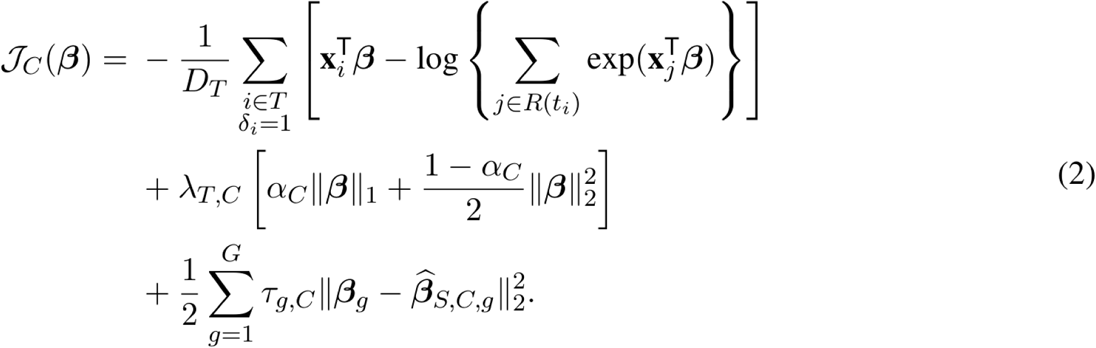

In both objectives, *λ* controls overall elastic-net regularization, *α* determines the L1/L2 mixture, and the layer-specific *τ* values control borrowing toward the source anchor. A larger *τ* pulls target coefficients within a layer closer to their source counterparts, favoring stronger borrowing when source and target associations are similar. When these associations differ, a smaller *τ* allows greater target-specific adaptation; *τ* = 0 removes direct anchoring for that layer.

For each candidate combination of screening budgets and tuning parameters, predictive loss is evaluated on the inner-validation fold and averaged across all five folds. Selection minimizes logistic log loss for binary prediction and negative Cox partial log-likelihood for time-to-event prediction. All methods use identical outer target splits, and performance is summarized as mean AUC across 10 repetitions (Fig. 1; Supplementary Fig. 1).

### 2.2 Simulation scenarios

We simulated binary outcomes with p = 500 predictors. Source, target-training, and target-test covariates were generated independently from a multivariate normal distribution with a common block-correlation structure: 50 groups of 10 predictors, correlation 0.50 within groups, and correlation 0.05 between groups. The source coefficient vector had 10 nonzero effects (three coefficients of 1.0, three of 0.7, and four of 0.4); binary outcomes were generated from a logistic model. Target coefficients were defined as *β*_T_ = *β*_S_ + *δ*, where each *δ*_j_ was drawn from N(0, *σ*_TL_^2^). We set *σ*_TL_^2^ to 0, 0.25, 1, or 4 to represent increasing source– target coefficient differences, each with increased source-target heterogeneity. Each setting used 300 source-training patients, 250 independent target-test patients, and a target-training size n_T_ of 60, 80, 100, 140, or 200. We generated 20 replicate datasets per setting and summarized target-test AUC by its mean and standard error.

### 2.3 TCGA-BRCA young patients survival prediction

We analyzed 523 female TCGA-BRCA primary-tumor data spanning six molecular layers: mRNA expression, miRNA expression, DNA methylation, copy-number variation, protein abundance, and somatic mutations. Molecular profiles were matched to age and survival records using TCGA sample identifiers (The Cancer Genome Atlas Network, 2012). Eligible patients had available clinical records and matched multi-omics profiles. Source cohorts comprised patients aged ≥60, ≥55, or ≥50 years, with patients aged <50 years forming the target cohort. After duplicate feature identifiers were removed, modality-specific preprocessing and quality filtering were followed by median imputation, feature-wise standardization, and layer-wise SIS. All data-dependent transformations and screening were fitted within the corresponding training partitions and applied unchanged to validation/test patients. Detailed preprocessing, filtering criteria, and screening procedures are provided in the Supplementary Methods and Fig. 1.

The endpoint was progression-free survival (PFI). For binary prediction at 2–4 years, patients were labeled alive at the horizon or dead before it; patients censored before the horizon were excluded from that horizon’s analysis. LSAT-EN was compared with Overall EN, fitted on pooled source and target-training data; Individual EN, fitted on target-training data only; glmtrans (Tian and Feng, 2022); and GAS-EN, an elastic-net adaptation of pooled fitting followed by target-specific correction. Cox analyses retained censored observations and compared LSAT-Cox-EN with Overall and Individual Cox-EN using Harrell’s C-statistic and 2–5-year IPCW AUC. BACC was used as a descriptive measure of horizon-specific classification performance, defining high risk by each model’s median held-out risk score within each split and excluding patients censored before the horizon with unknown event status. Proportional hazards were assessed using Schoenfeld residual tests in auxiliary Cox models relating held-out predicted risk scores to survival; diagnostic procedures are described in the Supplementary Methods (Grambsch and Therneau, 1994).

### 2.4 Risk stratification and restricted mean progression-free time analysis

LSAT-EN risk scores were calculated as *r*_i_(*t*) = 1 − P̂(*Y*_i_(*t*) = 1 | *X*_i_) for held-out target patients aged <50 years, where *X*_i_ denotes patient *i*’s preprocessed multi-omics feature vector and *Y*_i_(*t*) = 1 indicates remaining event-free beyond prediction horizon *t*; higher scores indicate greater predicted event risk. Patients were classified into low- and high-risk groups using the median risk score. Restricted mean event-free time was estimated from Kaplan–Meier curves through 2–5 years. Difference is calculated RMST_low-risk −RMST_high-risk such that positive values indicated longer restricted mean PFI in the predicted low-risk group. RMST differences were also calculated within molecular and clinical subgroups when sufficient samples were available. Confidence intervals for RMST differences were estimated using the unadjusted RMST difference output from the survRM2 package in R. (Uno et al., 2014)

### 2.5 Molecular Correlates of LSAT-EN–Derived Risk Scores

To assess whether LSAT-derived risk scores capture biologically interpretable molecular programs, we correlated held-out risk scores (calculated from 6 omics) for target patients aged <50 years with Hallmark pathway scores and miRNA features at 2–5-year PFI horizons (for the survival model). Hallmark gene sets were obtained from the MSigDB Hallmark collection (Liberzon et al., 2015), and pathway scores were calculated from z-scored mRNA expression values for genes available in each gene set. miRNA features were annotated using HMDD v4.0 to identify experimentally supported BC-associated miRNAs (Cui et al., 2023). Spearman correlations were calculated between LSAT risk scores and each Hallmark pathway score and miRNA feature, ranked correlation plots were used to visualize risk-associated pathway and miRNA patterns. Hallmark pathway scores and HMDD-annotated miRNA abundances were displayed using row-wise standardized values to visualize global molecular patterns across predicted risk groups.

## 3 Results

### 3.1 LSAT-EN improved prediction through layer-specific source borrowing

LSAT-EN incorporated six molecular layers: mRNA, miRNA, DNA methylation, copy-number variation, protein abundance, and somatic mutations. In each of ten repeated splits, 70% of target patients were used for training and 30% for testing. Within each inner training partition, modality-specific preprocessing and layer-wise SIS constructed candidate feature panels using source and target-training data, followed by source-only elastic-net fitting to estimate coefficient anchors. Inner fivefold cross-validation selected layer-specific screening budgets, elastic-net regularization parameters, and borrowing strengths. The selected pipeline was reconstructed using all source and target-training data, source anchors were re-estimated, and the target model was refitted with these anchors held fixed. Held-out predictions were used to evaluate performance and conduct downstream risk-stratification and molecular-correlation analyses (Fig. 1).

In group correlated data simulations, across target-training sizes n_T_ = 80, 100, 140, and 200, the group-tau method had the highest mean test AUC at every nonzero source–target coefficient-shift variance tested. Its greatest performance advantage occurs at highest source-target heterogeneity *σ*_TL_^2^ = 4, its AUC ranged from 0.689 to 0.789, compared with 0.561–0.592 for Overall EN, 0.629–0.747 for Individual EN, 0.590–0.721 for glmtrans EN, and 0.625–0.747 for GAS-EN. At *σ*_TL_^2^ = 0.25, the corresponding ranges were 0.747–0.803 for group-tau, 0.683–0.710 for Overall EN, 0.646–0.767 for Individual EN, 0.640–0.733 for glmtrans EN, and 0.735–0.767 for GAS-EN. At *σ*_TL_^2^ = 1, they were 0.683–0.788, 0.612–0.651, 0.634–0.766, 0.606–0.735, and 0.683–0.759, respectively. When there was no source–target coefficient shift (*σ*_TL_^2^ = 0), all methods performed well: group-tau achieved 0.943–0.957, Overall EN 0.954–0.961, Individual EN 0.925–0.954, glmtrans EN 0.956–0.962, and GAS-EN 0.945–0.961. glmtrans EN achieved the highest AUC when source and target coefficients were identical (*σ*_TL_^2^ = 0), with LSAT-EN performing comparably (Table 1; Supplementary Fig. 2). These results suggest that LSAT-EN maintains competitive performance when pooling is appropriate, while offering greater benefits under substantial source–target heterogeneity and limited target-training samples.

**Table 1.** Binary Transfer Simulation AUC Results.

| nT | Shift variance | glmtrans EN | Overall EN | Individual EN | GAS-EN | Group-tau EN |
|---|---|---|---|---|---|---|
| 200 | 0 | 96.2 (0.3) | 96.1 (0.3) | 95.4 (0.3) | 96.1 (0.3) | 95.7 (0.3) |
|  | 0.25 | 73.3 (1.5) | 71.0 (1.3) | 76.7 (1.0) | 76.7 (1.4) | 80.3 (0.8) |
|  | 1 | 73.5 (0.9) | 65.1 (1.4) | 76.6 (0.9) | 75.9 (1.5) | 78.8 (0.9) |
|  | 4 | 72.1 (1.2) | 59.2 (1.4) | 74.7 (1.6) | 74.7 (1.8) | 78.9 (0.9) |
| 140 | 0 | 95.8 (0.2) | 95.6 (0.2) | 94.5 (0.4) | 95.1 (0.2) | 95.2 (0.2) |
|  | 0.25 | 72.1 (1.4) | 68.3 (1.6) | 74.7 (1.1) | 74.6 (1.4) | 76.6 (1.1) |
|  | 1 | 65.9 (1.6) | 61.7 (1.1) | 69.7 (1.6) | 70.6 (1.2) | 73.7 (1.7) |
|  | 4 | 68.1 (1.3) | 56.1 (1.6) | 70.6 (1.5) | 73.3 (1.6) | 74.4 (1.5) |
| 100 | 0 | 96.2 (0.2) | 96.0 (0.2) | 93.3 (0.4) | 95.1 (0.4) | 95.1 (0.2) |
|  | 0.25 | 69.7 (2.1) | 69.9 (1.5) | 71.8 (1.8) | 75.5 (1.5) | 76.3 (1.4) |
|  | 1 | 65.3 (1.6) | 61.2 (1.2) | 65.8 (2.0) | 69.9 (1.7) | 70.3 (1.9) |
|  | 4 | 59.0 (1.8) | 56.1 (1.3) | 62.9 (2.0) | 62.5 (2.2) | 68.9 (1.6) |
| 80 | 0 | 95.6 (0.2) | 95.4 (0.2) | 92.5 (0.4) | 94.5 (0.3) | 94.3 (0.4) |
|  | 0.25 | 64.0 (2.3) | 70.4 (1.4) | 64.6 (2.0) | 73.5 (0.8) | 74.7 (1.0) |
|  | 1 | 60.6 (1.6) | 61.2 (1.5) | 63.4 (1.9) | 68.3 (1.6) | 68.3 (1.5) |
|  | 4 | 60.2 (1.6) | 56.2 (1.7) | 63.7 (1.8) | 64.6 (1.6) | 68.9 (1.2) |

In the binary survival model, LSAT-EN consistently outperformed non-TL and conventional TL methods across prediction horizons and source age groups, with patients aged <50 years forming the target cohort. LSAT-EN achieved 2-, 3-, and 4-year AUCs of 0.903, 0.813, and 0.819 with the ≥60-year source cohort (**Fig. 2A**); 0.906, 0.722, and 0.787 with the ≥55-year cohort (**Fig. 2B**); and 0.919, 0.770, and 0.737 with the ≥50-year cohort (**Fig. 2C**). Relative improvements ranged from 3.1%–22.7% over Overall EN, 4.8%– 39.3% over Individual EN, 9.2%–54.4% over glmtrans, and 4.9%–9.0% over GAS-EN. For time-to-event prediction, LSAT-Cox-EN achieved 2-, 3-, 4-, and 5-year IPCW AUCs of 0.945, 0.921, 0.874, and 0.901 with the ≥60-year source cohort (**Fig. 3A**); 0.936, 0.890, 0.879, and 0.887 with the ≥55-year cohort (**Fig. 3C**); and 0.855, 0.765, 0.809, and 0.781 with the ≥50-year cohort (**Fig. 3E**). Relative improvements ranged from 0.3%–22.7% over Overall Cox-EN and 1.1%–17.0% over Individual Cox-EN. Using descriptive median-risk classification, LSAT-Cox-EN achieved corresponding BACC values of 0.787, 0.816, 0.816, and 0.838 with the ≥60-year source cohort (**Fig. 3B**); 0.763, 0.772, 0.767, and 0.766 with the ≥55-year cohort (**Fig. 3D**); and 0.758, 0.697, 0.727, and 0.723 with the ≥50-year cohort (**Fig. 3F**). Across the 12 age–horizon settings, the mean relative improvement in BACC over Overall Cox-EN was 13.0%, while relative improvements over Individual Cox-EN ranged from 4.6%–14.3%. LSAT-Cox-EN also achieved the highest mean Harrell’s C-statistic across all three source-age cohorts, with values of 0.896, 0.862, and 0.786 for the ≥60-, ≥55-, and ≥50-year cohorts, respectively (Fig. 3G).

**Figure 2.**
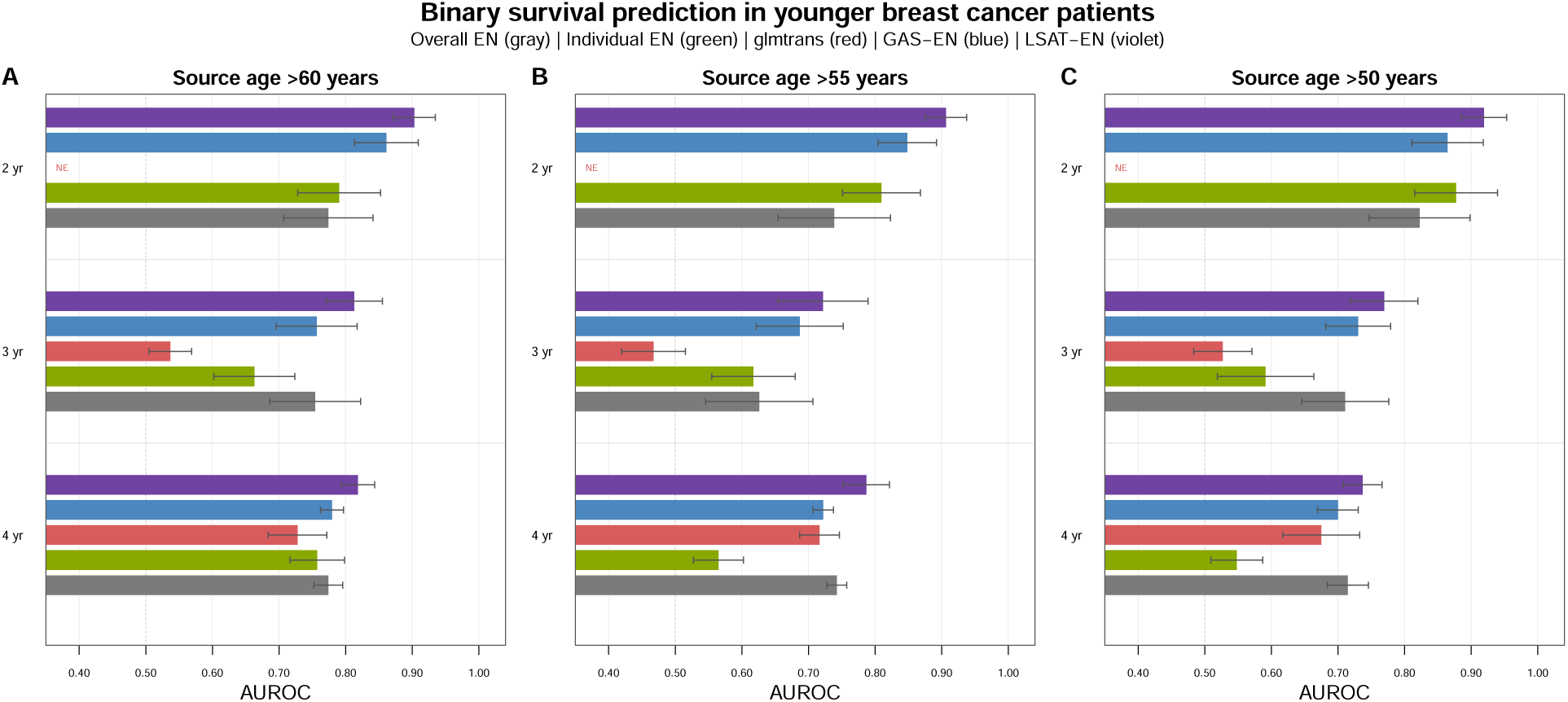
Binary survival prediction in younger breast cancer patients. Bars show mean held-out target-cohort AUROC for 2-, 3-, and 4-year overall survival prediction using selected multi-omics feature panels. Target patients were aged <50 years; source patients were aged >60 (A), >55 (B), and >50 years (C). Methods include Overall EN (gray), Individual EN (green), glmtrans (red), GAS-EN (blue), and LSAT-EN (violet). Error bars indicate descriptive standard errors across 10 repeated target train/test splits. NE denotes a nonevaluable result; glmtrans had no evaluable 2-year results. LSAT-EN achieved the highest mean AUROC in all nine age–horizon settings.

**Figure 3.**
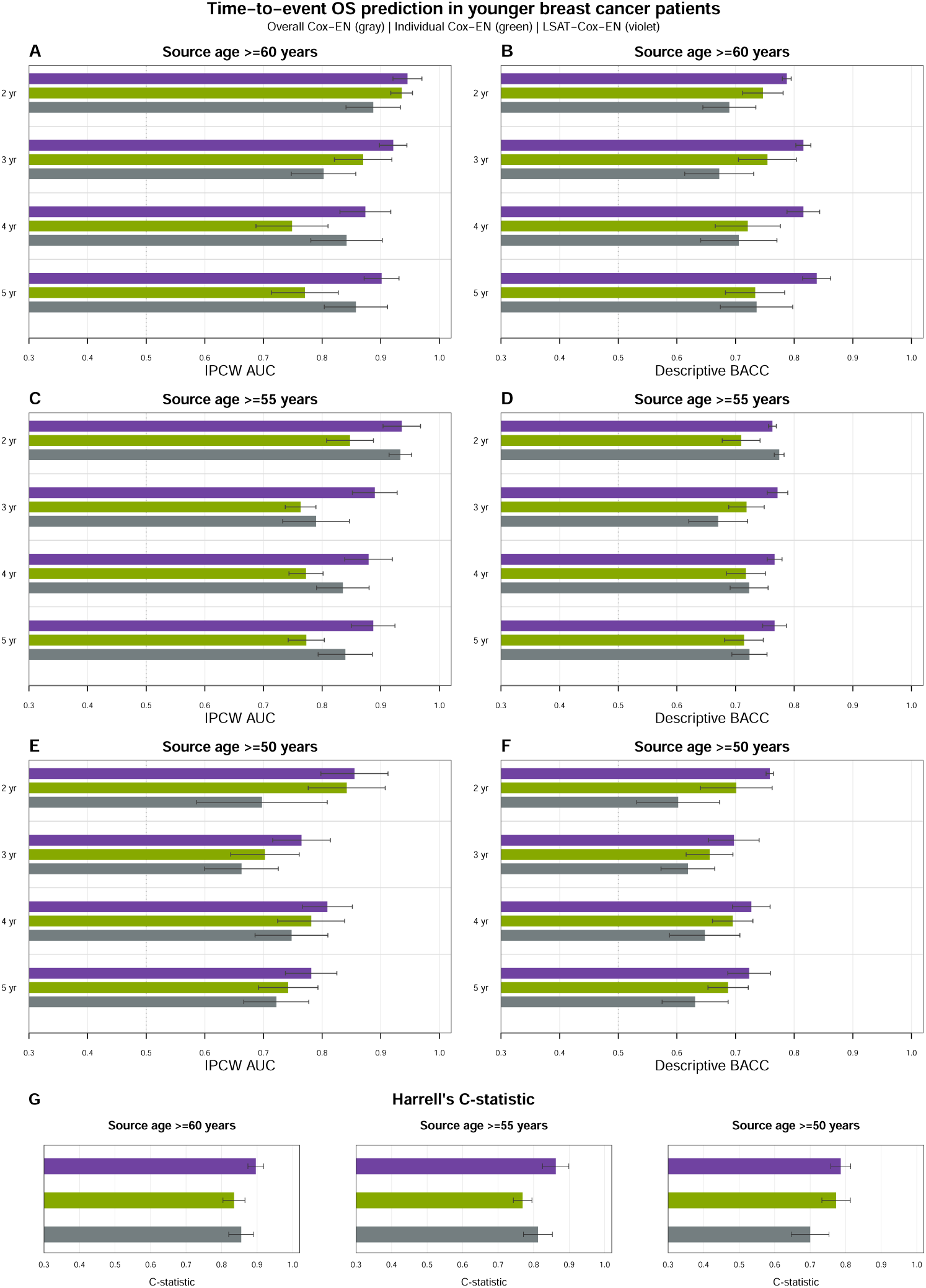
Time-to-event overall survival prediction in younger breast cancer patients. Bars show mean held-out target-cohort IPCW AUC (A, C, E) and descriptive balanced accuracy (BACC; B, D, F) at 2–5 years, and Harrell’s C-statistic for the three source-age cohorts (G). Target patients were aged <50 years; source patients were aged ≥60 (A, B), ≥55 (C, D), and ≥50 years (E, F). Methods include Overall Cox-EN (gray), Individual Cox-EN (green), and LSAT-Cox-EN (violet). BACC used each method’s median held-out risk score within each split to define high- and low-risk predictions, excluding patients censored before the evaluation horizon. Error bars indicate standard errors across 10 repeated target train/test splits.

### 3.2 LSAT-EN maintained robust PFI prediction and risk stratification across BC molecular and clinical subtypes

To examine whether LSAT-derived risk groups were clinically interpretable, we compared PFI event composition and restricted mean survival time (RMST) differences between LSAT low-risk and high-risk groups (Fig. 4), with risk scores calculated as one minus the predicted probability of remaining event-free beyond each prediction horizon. RMST estimates the expected progressionfree survival time up to a specified PFI time and provides an absolute time-scale summary that can be easier to interpret clinically than a hazard ratio, and has the advantage of not relying on underlying model assumptions. At the 4- and 5-year PFI prediction, the low-risk group included 72 samples, and the high-risk group included 73 samples, with PFI events less frequent in the low-risk group than in the high-risk group at 4 years approximately 3% versus 11%, respectively, and at 5 years, 4% versus 15% respectively (Fig. 4E, G). Similar patterns were observed at 2 and 3 years, with observed PFI events in 0% versus approximately 5% and 0% versus 8% of low- and high-risk patients, respectively (Fig. 4A, C), although early censoring left some patients’ event status unknown at each horizon. These analyses supported the robustness of LSAT - derived risk stratification across molecular and clinical subgroups. LSAT low-risk patients generally had longer mean PFI than predicted high-risk patients, with positive RMST differences observed across most PAM50, stage, tumor, lymph node, and metastatic subgroups (Fig. 4B, D, F, H). The overall RMST difference remained positive across all PFI time settings, and the direction of separation was maintained in many individual subgroups despite variability in effect size and confidence intervals. These results indicate that LSAT-derived risk groups captured risk differences across PFI time settings beyond the overall cohort and across heterogeneous subtypes. Overall, these findings support the potential clinical utility of LSAT for risk stratification in heterogeneous BC patients.

**Fig 4.**
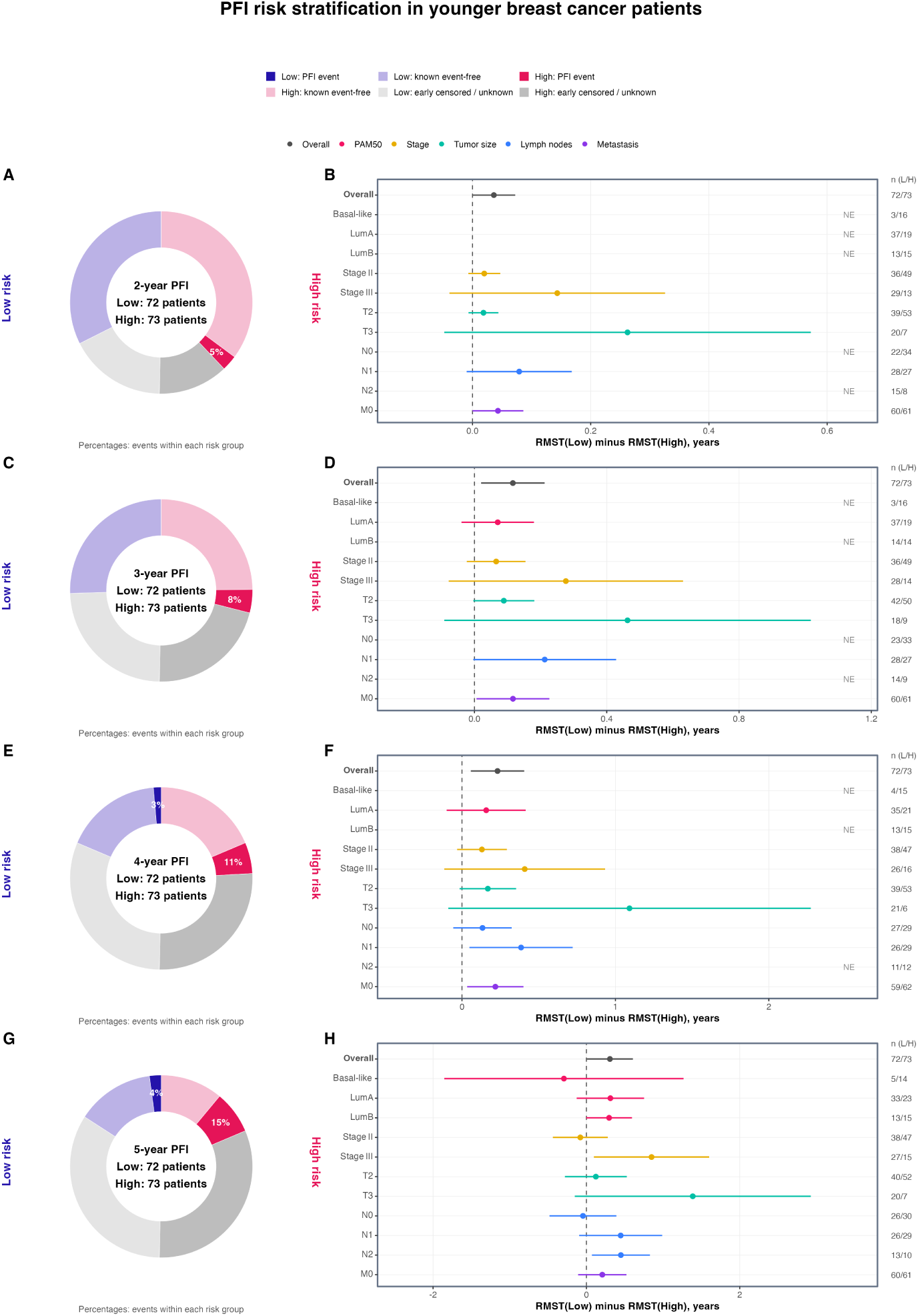
LSAT-derived risk groups differed in PFI event burden and restricted mean survival time. LSAT low- and high-risk groups were evaluated at 2 (**A, B**), 3 (**C, D**), 4 (**E, F**), and 5 years (**G, H**). Donut charts show PFI event and non-event composition; centers indicate sample size. Risk groups were defined by the median LSAT risk score at each PFI time setting. Forest plots show RMST differences (RMST_low − RMST_high) for the overall cohort and eligible subgroups. Positive values indicate longer restricted mean PFI in the low-risk group. Points indicate RMST differences; horizontal lines indicate 95% confidence intervals.

### 3.3 LSAT-EN risk scores captured breast cancer-relevant molecular programs across multi-omics data

To investigate whether LSAT-derived risk scores capture biologically interpretable signals, we calculated correlations between risk scores and molecular features, including MSigDB Hallmark pathway score and miRNA abundance across PFI time settings (**Fig. 5**). This analysis was designed to determine whether MOTLAB-derived risk scores in younger patients were associated with BC-related molecular pathways. We used the Hallmark gene set collection from MSigDB because it summarizes redundant gene sets into coherent biological states and processes such as proliferation, immune response, DNA damage, metabolism, and oncogenic signaling (Liberzon et al., 2015). In parallel, we annotated miRNAs using the Human MicroRNA Disease Database (HMDD v4.0), a curated resource of experimentally supported associations between human miRNAs and diseases, to identify miRNAs linked to BC (Cui et al., 2023).

**Fig. 5.**
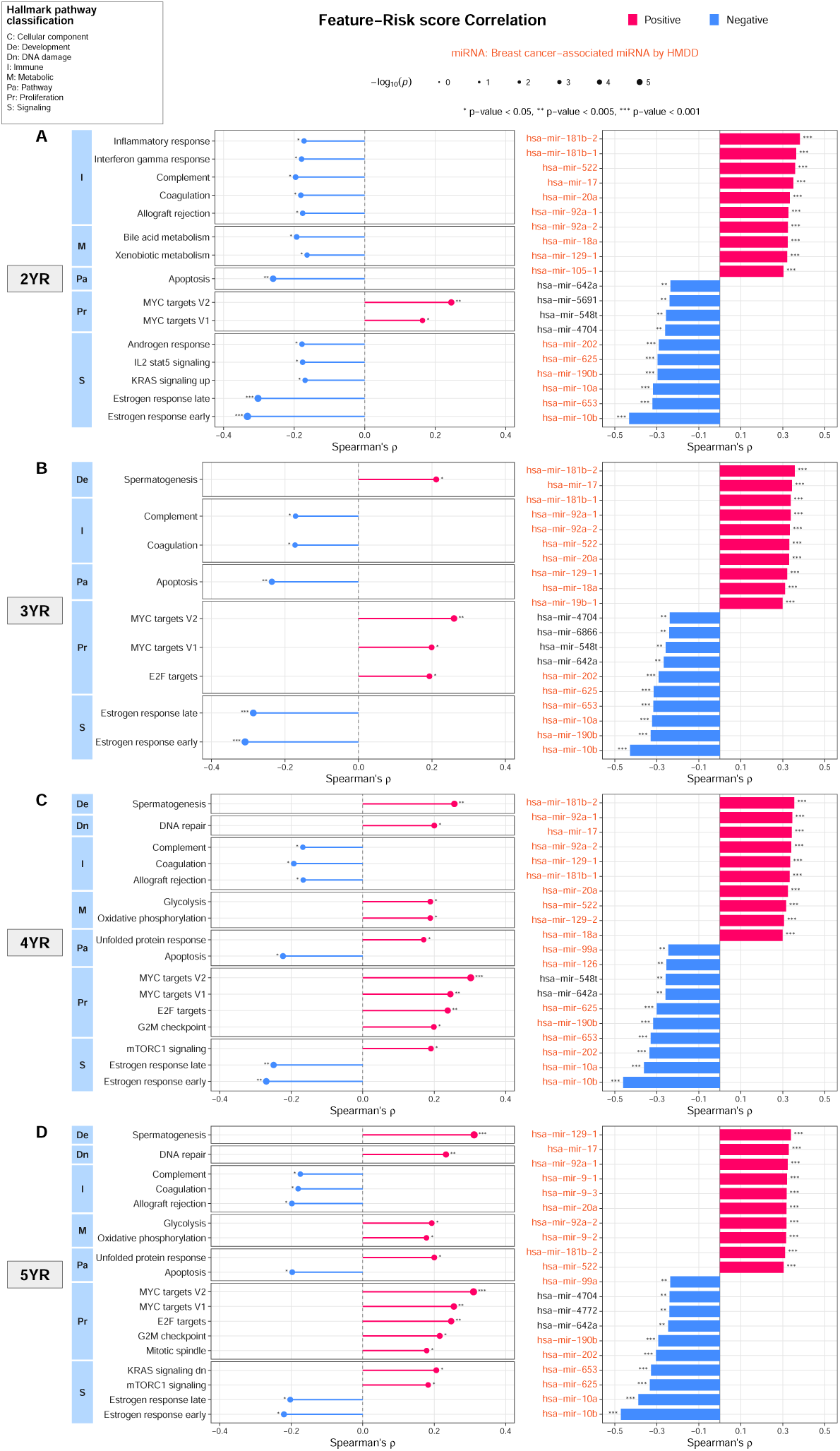
LSAT-EN risk scores were associated with Hallmark pathway and miRNA programs. Spearman correlations between MOTLAB risk scores and Hallmark pathway scores or miRNA abundance at 2 (A), 3 (B), 4 (C), and 5 years (D). Left panels show Hallmark pathways grouped by process category; right panels show risk-associated miRNAs. Red and blue indicate positive and negative correlations with LSAT risk, respectively. Asterisks indicate nominal Spearman p values (*p < 0.05, **p < 0.005, ***p < 0.001). Orange labels indicate BC-associated miRNAs identified from HMDD v4 disease entries containing “breast.”

At the 5-year PFI prediction, higher LSAT-EN risk was associated with several Hallmar pathways linked to aggressive BC, including mTORC1 signaling (Loi et al., 2010), DNA replication (Shetty et al., 2005), E2F targets, G2M checkpoint, Mitotic Spindle, MYC targets V1/V2, glycolysis (Zheng et al., 2022), and oxidative phosphorylation (El-Botty et al., 2023) (Fig. 5D). BC studies have linked increased MYC target activity (Schulze et al., 2020), ROS abundance (Oshi et al., 2022a), and oxidative phosphorylation-associated gene expression (El-Botty et al., 2023) to aggressive BC or poor survival in BC, while mTORC1 signaling (Loi et al., 2010) and glycolysis-related signatures (Zheng et al., 2022) have also shown subtype-specific prognostic associations. These findings support the biological relevance of the molecular variation captured by LSAT risk. In contrast, pathways associated with lower risk score (pathways negatively associated with risk scores) include immune pathways like complement, coagulation, allograft rejection, as well as estrogen response and apoptosis (Fig 5D), which is consistent with the favorable prognostic associations of antitumor immune activity in some breast cancer subtypes (Ali et al., 2016). Across the 2– 5-year PFI prediction horizons, higher LSAT risk was associated primarily with metabolic and tumorrelated pathways at later horizons, particularly 5 years, whereas lower LSAT risk was associated with immune-related pathways at earlier horizon (e.g. 2-years) (Fig. 5B–D). For example, 2 year pathways negatively associated with risk scores in addition are inflammatory, interferon gamma response, androgen, and IL2 stat5 signaling. This pattern is consistent with subtype-specific evidence linking type I interferon activity to favorable outcomes (Cao et al., 2023), IFN-γ signaling to prognosis (Heimes et al., 2020), and enhanced tumor immune activity to improved survival in TNBC and HER2-positive BC(Denkert et al., 2018; Oshi et al., 2022b), consistent with evidence linking greater antitumor immune activity and interferon-related transcriptional pathways to better outcomes in BC. MYC targets V1 and V2 correlated with risk scores from all three PFI predictions, while more pathways become significant at 4-5 years, including glycolysis, oxidative phosphorylation (metabolic events), G2M checkpoint and mitotic spindle (proliferation), and important KRAS and mTOR signaling pathways. Conversely, complement, coagulation, and allograft-rejection showed recurrent inverse associations with LSAT risk.

Consistent with findings from pathway analysis, HMDD-annotated BC-associated miRNAs continued to appear among the strongest positive and negative risk correlations at these PFI time settings (labeled in red). At 2-years, higher predicted PFI risk was positively correlated with HMDD-annotated miRNAs including hsa-miR-181b-2, hsa-miR-522, hsa-miR-17, hsa-miR-20a, and hsa-miR-92a-1/2 (Spearman’s ρ = 0.32–0.38), whereas hsa-miR-202, hsa-miR-625, hsa-miR-190b, hsa-miR-10a, and hsa-miR-10b were negatively correlated with risk (ρ = −0.43 to −0.29). Prior BC studies reported tumor-suppressive activity for miR-625 (Zhou et al., 2016) and longer metastasis-free and event-free survival with higher miR-190b expression (Cizeron-Clairac et al., 2015). Compared with the 2-year results, 5-year associations additionally included hsa-miR-9-1/2/3 positively (ρ = 0.316–0.319) and hsa-miR-99a and hsa-miR-4772 negatively (ρ = −0.236 and −0.240, respectively). For example, miR-9 promotes breast cancer invasiveness and metastasis through E-cadherin suppression, whereas miR-99a inhibits tumor growth through mTOR targeting (Ma et al., 2010; Hu et al., 2014). Remarkably, the most positive and negative correlation were breast cancer miRNA, across 2-5 years the 10 most positive correlations and 6 most negative correlations were HMDD-annotated BC miRNAs. These results suggest that LSAT-derived risk scores capture BC-relevant post-transcriptional regulatory variation. Overall, these findings suggest that LSAT-derived risk scores that remained informative across molecular and clinical subgroups and were associated with BC-relevant pathway and miRNA patterns, supporting their clinical and biological interpretability.

## 4 Discussion

In this study, we proposed LSAT-EN, a weighted multi-modal transfer learning framework with variable selection. LAST-EN demonstrated that weighted multi-omics integration with transfer learning (TL) and variable selection by EN provides a principled strategy to improve model performance and mitigate age disparities in breast cancer (BC). By integrating all omics, LSAT-EN leverages a richer representation of tumor biology. This aligns with studies showing that multi-omics integration improves cancer subtyping and survival prediction by capturing complementary molecular information (Wang et al., 2014; Ramazzotti et al., 2018; Shen et al., 2012; Chaudhary et al., 2018). In addition, optimization of weight combinations for omics integration allowed LSAT-EN to emphasize the most informative omics data rather than assuming equal contributions from each omics data. This aligns with recent methodological advances that highlight the advantages of weighted integration for cancer subtyping and prognosis (Cao et al., 2024; Wei et al., 2022; Ramazzotti et al., 2018; Duan et al., 2024).

LSAT-EN’s advantage under source–target heterogeneity, particularly with small target samples, supports the value of its layer-specific *τ* penalties in balancing source information with target-specific adaptation. This tunable borrowing mechanism distinguishes LSAT-EN from the glmtrans and GAS-EN formulations evaluated here and extends to both binary and Cox models. When coefficients were identical, methods performed similarly, with glmtrans achieving the highest AUC. Beyond prediction, unlike MOTL (Hirst et al., 2025), which transfers multi-omics latent factors, and MOTLAB (Baek et al., 2025), which uses similarity-weighted integration and neural-network transfer, LSAT-EN retains explicit feature-level coefficients. Its elastic-net regularization accommodates correlated predictors and performs variable selection, supporting downstream biological pathway analyses and risk stratification.

The molecular analyses support the biological relevance of the risk information learned by LSAT-EN. LSAT-derived risk scores were associated with established MSigDB Hallmark pathways, breast cancer-associated miRNAs curated in HMDD, linking model-derived risk to transcriptional, post-transcriptional, variation relevant to breast cancer biology (The Cancer Genome Atlas Network, 2012; Liberzon et al., 2015; Cui et al., 2023). This multi-omics context is relevant given previous evidence of molecular heterogeneity in BC and ancestry-associated variation across transcriptomic and post-transcriptomic features in TCGA cohorts(Huo et al., 2017; Carrot-Zhang et al., 2020). However, previous ancestry-based molecular analyses have also highlighted the importance of tumor subtype and technical factors when interpreting such differences (Carrot-Zhang et al., 2020). Accordingly, these findings provide biological context for the predictive signals and support the interpretation of the identified pathways and miRNAs features as molecular correlates of PFI. The consistently improved predictive performance across molecular and clinical subgroups supports the robust applicability of LSAT-EN across diverse BC characteristics. With these findings, LSAT-EN provide a generalizable framework for improving personalized prediction in BC data-minority groups.

To improve the method’s robustness and effectiveness, several additional considerations should be addressed in future studies aimed at reducing BC disparities. The analysis was based on a single TCGA cohort with a relatively small younger age group, and even fewer target events, especially at earlier timepoints (e.g. 2-3 years) and therefore requires validation in independent, diverse larger multi-institutional cohorts or cancer registries. The method could also be extended to other disparities or minorities, e.g. sex or ethnicity, and include molecular, clinical and treatment features apart from multi-omics data.

## 5 Conclusion

We developed LSAT-EN, a layer-weighted multi-omics transfer-learning framework that adaptively borrows source information to improve prediction in data-limited target cohorts. By combining target-specific estimation with elastic-net variable selection, LSAT-EN supports both prognostic modeling and molecular-based risk stratification. Its application to younger breast cancer patients illustrates its potential to advance cancer machine learning for broader underrepresented populations and contribute to efforts to reduce cancer disparities.

## Supplementary Material

Supplementary tables and figures are provided in a separate supplementary file.

## Data availability

The multi-omics and clinical outcome data used in this study were derived from The Cancer Genome Atlas breast invasive carcinoma cohort. TCGA molecular and clinical resources are available through the National Cancer Institute Genomic Data Commons (https://portal.gdc.cancer.gov/). LSAT-EN is available at https://github.com/wguo3773/star-en.

