## Supplementary Materials for "Layer-Weighted Source-Anchored Transfer Learning for Multi-Omics Survival Prediction and Risk Stratification in Younger Breast Cancer Patients"

#### Supplementary Methods

##### Multi Omics Preprocessing and Feature Screening

**Sample matching and cohort definition.** TCGA-BRCA primary-tumor molecular profiles were matched to age and survival records using TCGA identifiers. Analyses included female patients with available age, endpoint information, and matched multi-omics profiles. Source cohorts were defined separately using age cutoffs of ≥60, ≥55, and ≥50 years; patients aged <50 years formed the target cohort. Duplicate feature identifiers were removed before analysis.

**Modality-specific preprocessing and filtering.** Six molecular layers were processed separately using the criteria below. Expression-detection thresholds were assessed on the CPM scale before log transformation; data-dependent filtering statistics were calculated from the corresponding training partition.

| Molecular layer | Preprocessing | Feature-retention criteria |
|---|---|---|
| mRNA | Convert raw counts to CPM; transform using $\log_2(\mathrm{CPM} + 1)$. | CPM >1 in ≥50% of training patients. |
| miRNA | Convert raw counts to CPM; transform using $\log_2(\mathrm{CPM} + 1)$. | CPM >0.1 in ≥30% of training patients. |
| DNA methylation | Use quality-controlled β values summarized at the gene level. | Observed in ≥30% of training patients; retain the top 50% by training-data IQR. |
| Copy-number variation | Use neutral-centered gene-level copy-number values. | Observed in ≥70% of training patients and training-data IQR >0. |
| Protein abundance | Use normalized reverse-phase protein array measurements. | Observed in ≥70% of training patients and training-data IQR >0. |
| Somatic mutations | Encode nonsynonymous mutations as binary gene-level indicators. | At least five training carriers; remove constant features. |

**Imputation and standardization.** Following quality filtering, missing predictor values were replaced with feature-specific training medians. Each retained feature was standardized by subtracting its training mean and dividing by its training standard deviation. Source and target predictors used the same fitted transformations to maintain compatible coefficient scales. Features with zero training variance were excluded.

**Layer-wise screening and budget selection.** Sure independence screening (SIS) ranked eligible features separately within each molecular layer using source and target-training outcomes. Binary prediction used an outcome-appropriate marginal logistic screening statistic, whereas time-to-event prediction used a marginal Cox screening statistic incorporating censoring. Candidate feature budgets were evaluated separately by layer:

| Molecular layer | Candidate retained-feature counts |
|---|---|
| mRNA | 250, 500, 1,000 |
| miRNA | 50, 100, 200 |
| DNA methylation | 250, 500, 1,000 |
| Copy-number variation | 100, 200, 400 |
| Protein abundance | 50, 100, all eligible features |
| Somatic mutations | 25, 50, 100 |

For each candidate budget, the highest-ranked eligible features were retained; when fewer features were available, all eligible features were retained. Selected features were concatenated across layers while preserving their layer labels. Budget combinations were selected jointly with the regularization parameters using inner cross-validation performance.

**Training-only implementation and evaluation.** Within each repeated 70% target-training/30% target-testing split, the target-training subset was divided into five stratified folds. For each inner iteration, filtering, imputation, standardization, and SIS were fitted using all source patients and four target-training folds, then applied unchanged to the remaining validation fold. Source-only elastic-net coefficient anchors were re-estimated whenever the selected panel or preprocessing scale changed and held fixed during the corresponding target-model fit. Mean validation logistic log loss or negative Cox partial log-likelihood selected the layer-specific budgets and parameters $\lambda$, $\alpha$, and $\tau_g$. The selected pipeline was reconstructed using all source and target-training patients, source anchors were refitted, and the target model was fitted using the target likelihood and fixed anchors. Performance was evaluated on untouched target-test patients and averaged across 10 repeated splits (Supplementary Fig. 1).

### Supplementary Figures and Tables

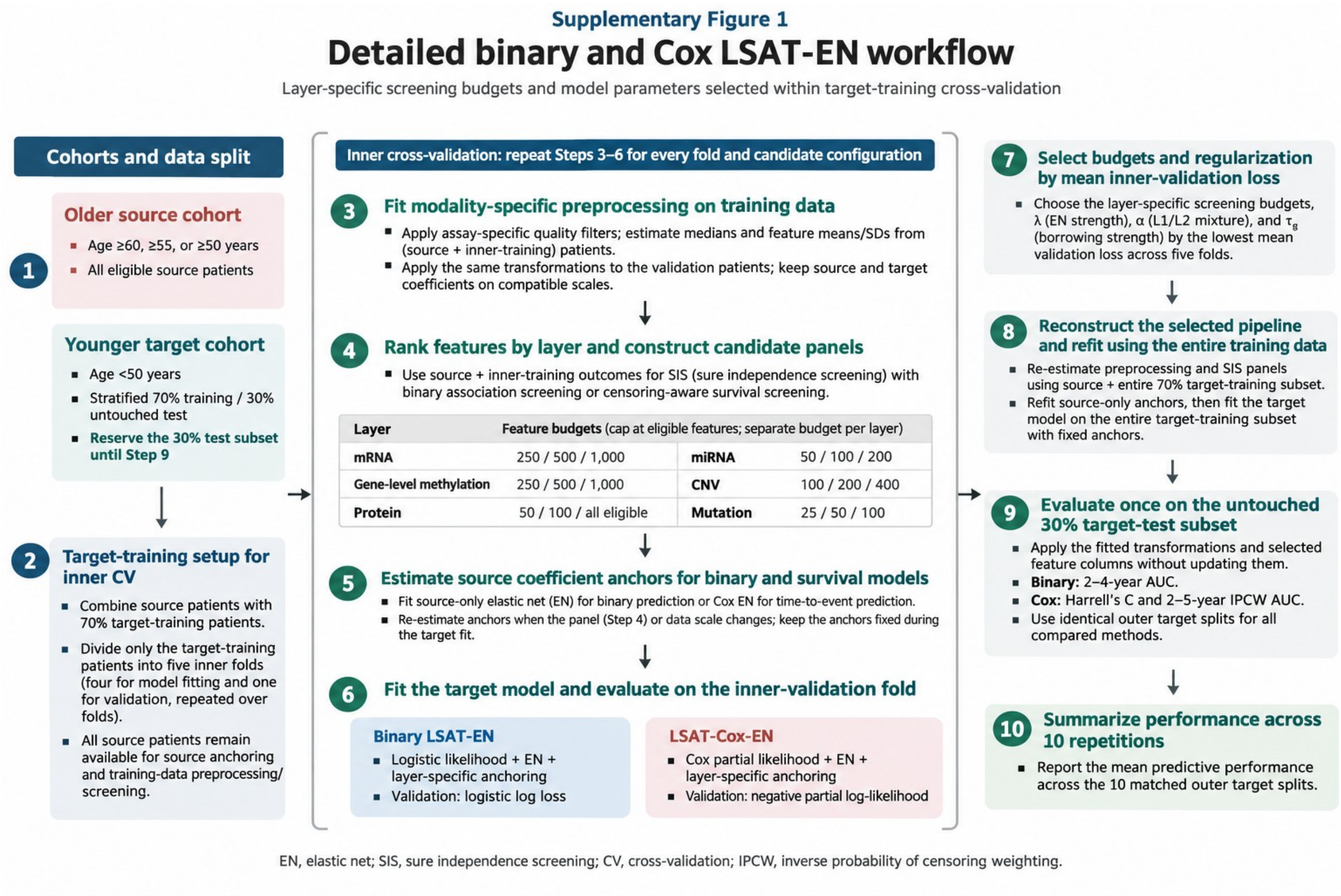


Supplementary Figure 1. Detailed training workflow for binary LSAT-EN and LSAT-Cox-EN. (1–2) Target patients aged <50 years undergo stratified 70% training/30% testing splits, with five-fold cross-validation within the target-training subset. Older source cohorts are defined using age cutoffs of ≥60, ≥55, or ≥50 years. (3–6) Within each cross-validation round, modality-specific filtering, imputation, standardization, and layer-wise SIS are fitted using source patients and four target-training folds to construct candidate feature panels; the remaining fold is reserved for validation. Source-only elastic-net models provide endpoint-specific coefficient anchors, re-estimated when the panel or preprocessing scale changes and held fixed during the corresponding target fit. (7) Mean validation loss selects layer-specific screening budgets and regularization parameters $\lambda, \alpha, \tau_g$, using logistic log loss for binary prediction and negative Cox partial log-likelihood for censored outcomes using source + full training data (8–9) The selected pipeline is reconstructed on the full training data and evaluated on untouched target-test patients. (10) Predictive performances are summarized across 10 matched outer splits. Binary evaluation uses 2–4-year AUC; Cox evaluation uses Harrell's C-statistic and 2–5-year IPCW AUC. Feature panels and selected budgets may vary across repetitions.

#### Supplementary Figure 2

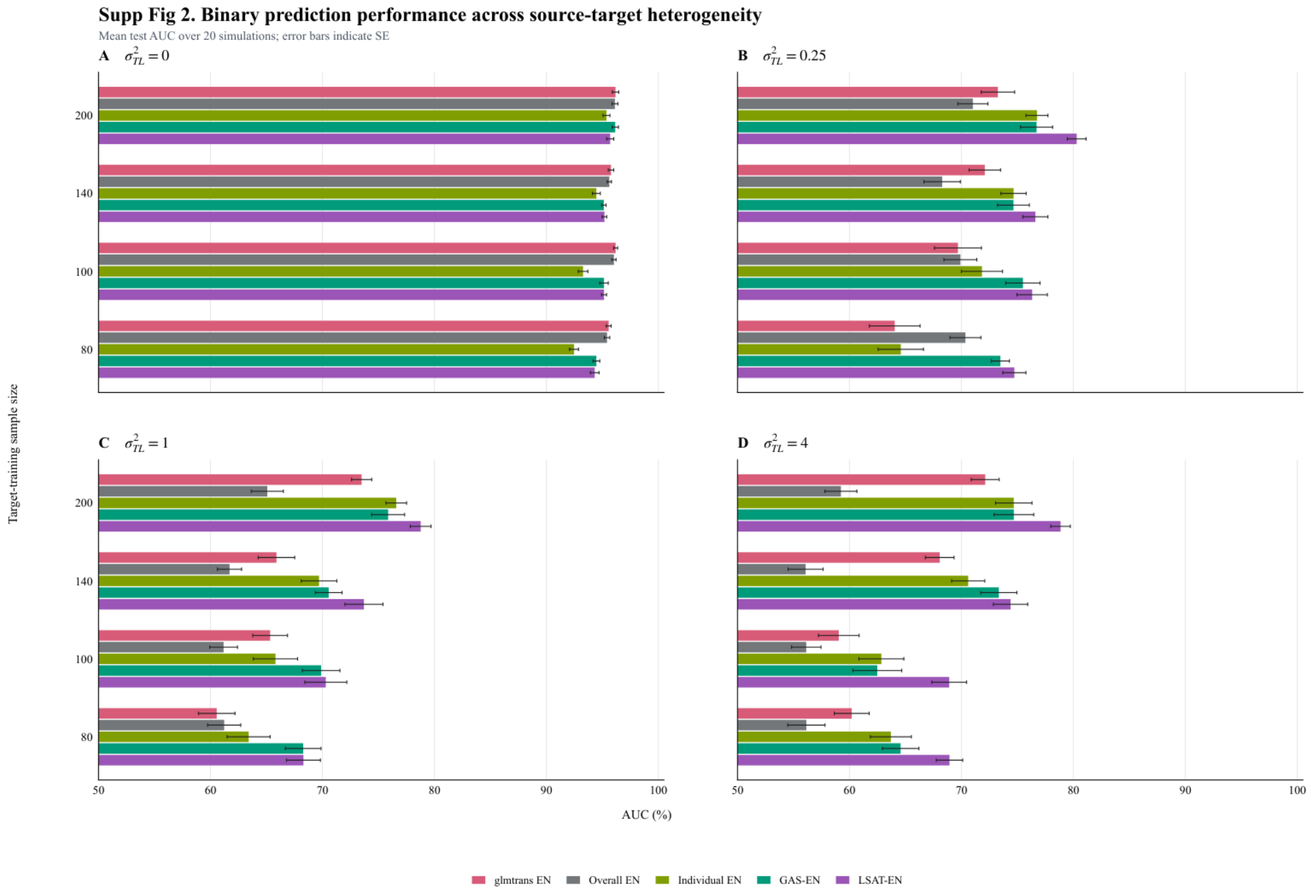


Supplementary Figure 2. Binary prediction performance across source–target heterogeneity and target-training sample sizes. Data is from Table 1. Panels A–D show source–target coefficient-shift variances of σ²TL = 0, 0.25, 1, and 4, respectively. Five methods are compared at target-training sample sizes of 80, 100, 140, and 200: glmtrans EN, Overall EN, Individual EN, GAS-EN, and LSAT-EN. Bars show mean out-of-sample AUC (%) across 20 simulations per setting; error bars represent SE. Each test set contains 250 observations. Higher AUC indicates better discrimination.

#### Supplementary Table 1

#### Binary survival prediction in the younger target cohort

Mean AUC (%) with SE in parentheses for the nine selected panels in Figure 2. The target cohort comprises patients aged <50 years. Bold indicates the highest mean AUC among evaluable methods in each row.

| Panel | Source age | Horizon (years) | Overall EN | Individual EN | glmtrans | GAS-EN | LSAT-EN |
|---|---|---|---|---|---|---|---|
| A | >60 | 2 | 77.4 (6.7) | 79.0 (6.2) | NA | 86.1 (4.8) | **90.3 (3.2)** |
| B | >60 | 3 | 75.4 (6.8) | 66.3 (6.1) | 53.7 (3.2) | 75.7 (6.1) | **81.3 (4.2)** |
| C | >60 | 4 | 77.4 (2.2) | 75.7 (4.1) | 72.8 (4.4) | 78.0 (1.7) | **81.9 (2.5)** |
| D | >55 | 2 | 73.9 (8.5) | 81.0 (5.8) | NA | 84.8 (4.4) | **90.6 (3.1)** |
| E | >55 | 3 | 62.6 (8.0) | 61.7 (6.3) | 46.7 (4.8) | 68.7 (6.5) | **72.2 (6.8)** |
| F | >55 | 4 | 74.3 (1.5) | 56.5 (3.8) | 71.7 (3.0) | 72.2 (1.5) | **78.7 (3.5)** |
| G | >50 | 2 | 82.3 (7.6) | 87.7 (6.2) | NA | 86.5 (5.4) | **91.9 (3.4)** |
| H | >50 | 3 | 71.1 (6.5) | 59.1 (7.2) | 52.7 (4.4) | 73.0 (4.9) | **77.0 (5.1)** |
| I | >50 | 4 | 71.5 (3.1) | 54.8 (3.9) | 67.5 (5.8) | 70.0 (3.1) | **73.7 (2.9)** |

Notes. SE = standard error across 10 repeated target train/test splits. NA indicates that glmtrans did not yield an evaluable 2-year result in panels A, D, and G.

#### Supplementary Table 2

#### Time to event prediction in the younger target cohort

Mean C-statistic, IPCW AUC, or descriptive balanced accuracy (BACC), with SE in parentheses. Current Figure 3 AUC panels are A, C, and E; BACC panels are B, D, and F. C-statistics are retained as additional results. The target cohort comprises patients aged <50 years. Bold indicates the highest mean in each row.

| Panel | Source age | Metric | Overall Cox-EN | Individual Cox-EN | LSAT-Cox-EN |
|---|---|---|---|---|---|
| - | ≥60 | C-statistic | 0.855 (0.035) | 0.835 (0.031) | **0.896 (0.022)** |
| A | ≥60 | 2-year IPCW AUC | 0.887 (0.046) | 0.936 (0.019) | **0.945 (0.025)** |
| A | ≥60 | 3-year IPCW AUC | 0.802 (0.055) | 0.870 (0.049) | **0.921 (0.023)** |
| A | ≥60 | 4-year IPCW AUC | 0.842 (0.061) | 0.749 (0.061) | **0.874 (0.043)** |
| A | ≥60 | 5-year IPCW AUC | 0.858 (0.054) | 0.771 (0.057) | **0.901 (0.030)** |
| B | ≥60 | 2-year descriptive BACC | 0.689 (0.045) | 0.747 (0.034) | **0.787 (0.008)** |
| B | ≥60 | 3-year descriptive BACC | 0.672 (0.059) | 0.754 (0.049) | **0.816 (0.013)** |
| B | ≥60 | 4-year descriptive BACC | 0.706 (0.065) | 0.721 (0.056) | **0.816 (0.028)** |
| B | ≥60 | 5-year descriptive BACC | 0.736 (0.062) | 0.733 (0.050) | **0.838 (0.024)** |
| - | ≥55 | C-statistic | 0.812 (0.041) | 0.769 (0.026) | **0.862 (0.037)** |
| C | ≥55 | 2-year IPCW AUC | 0.933 (0.019) | 0.847 (0.040) | **0.936 (0.032)** |
| C | ≥55 | 3-year IPCW AUC | 0.789 (0.057) | 0.763 (0.026) | **0.890 (0.038)** |
| C | ≥55 | 4-year IPCW AUC | 0.835 (0.045) | 0.772 (0.029) | **0.879 (0.040)** |
| C | ≥55 | 5-year IPCW AUC | 0.839 (0.046) | 0.773 (0.031) | **0.887 (0.037)** |
| D | ≥55 | 2-year descriptive BACC | **0.775 (0.008)** | 0.710 (0.032) | 0.763 (0.007) |
| D | ≥55 | 3-year descriptive BACC | 0.670 (0.050) | 0.719 (0.030) | **0.772 (0.018)** |
| D | ≥55 | 4-year descriptive BACC | 0.723 (0.032) | 0.718 (0.033) | **0.767 (0.013)** |
| D | ≥55 | 5-year descriptive BACC | 0.724 (0.030) | 0.714 (0.033) | **0.766 (0.020)** |
| - | ≥50 | C-statistic | 0.700 (0.053) | 0.773 (0.040) | **0.786 (0.028)** |
| E | ≥50 | 2-year IPCW AUC | 0.697 (0.111) | 0.842 (0.066) | **0.855 (0.057)** |
| E | ≥50 | 3-year IPCW AUC | 0.662 (0.063) | 0.702 (0.058) | **0.765 (0.049)** |
| E | ≥50 | 4-year IPCW AUC | 0.748 (0.062) | 0.781 (0.057) | **0.809 (0.043)** |
| E | ≥50 | 5-year IPCW AUC | 0.722 (0.055) | 0.742 (0.051) | **0.781 (0.044)** |
| F | ≥50 | 2-year descriptive BACC | 0.602 (0.071) | 0.701 (0.061) | **0.758 (0.006)** |
| F | ≥50 | 3-year descriptive BACC | 0.619 (0.046) | 0.656 (0.040) | **0.697 (0.043)** |
| F | ≥50 | 4-year descriptive BACC | 0.647 (0.060) | 0.695 (0.035) | **0.727 (0.032)** |
| F | ≥50 | 5-year descriptive BACC | 0.631 (0.056) | 0.687 (0.034) | **0.723 (0.036)** |

Notes. SE = standard error across evaluable repeated target train/test splits. BACC uses each model's held-out median risk within each split; patients censored before the horizon with unknown status were excluded. SEs across overlapping splits are descriptive. IPCW = inverse probability of censoring weighting.
